# Who gets lost and why: A representative cross-sectional survey on sociodemographic and vestibular determinants of wayfinding strategies

**DOI:** 10.1101/419986

**Authors:** Susanne Ulrich, Eva Grill, Virginia L. Flanagin

**Author notes:** corresponding author (VLF).

## Abstract

When we think of our family and friends, we probably know someone who is good at finding their way and someone else that easily get lost. We still know little about the biological and environmental factors that influence our navigational ability. Here, we investigated the frequency and sociodemographic determinants of wayfinding and their association with vestibular function in a representative cross-sectional sample (N = 783) of the adult German-speaking population. Wayfinding was assessed using the Wayfinding Strategy Scale, a self-report scale that produces two scores for each participant representing to what degree they rely on route-based or orientation (map-based) strategies. We were interested in the following research questions: (1) the frequency and determinants of wayfinding strategies in a population-based representative sample, (2) the relationship between vestibular function and strategy choice and (3) how sociodemographic factors influence general wayfinding ability as measured using a combined score from both strategy scores. Our linear regression models showed that being male, having a higher education, higher age and lower regional urbanization increased orientation strategy scores. Vertigo/dizziness reduced the scores of both the orientation and the route strategies. Using a novel approach, we grouped participants by their combined strategy scores in a multinomial regression model, to see whether individuals prefer one strategy over the other. The majority of individuals reported using either both or no strategy, instead of preferring one strategy over the other. Young age and reduced vestibular function were indicative of using no strategy. In summary, wayfinding ability depends on both biological and environmental factors; all sociodemographic factors except income. Over a third of the population, predominantly under the age of 35, does not successfully use either strategy. This represents a change in our wayfinding skills, which may result from the technological advances in navigational aids over the last few decades.

## Introduction

Wayfinding; the ability to find one’s way in an unfamiliar environment, has an outstanding place among our cultural skills. The ability to find one’s way in a complex environment is no trivial feat. It is therefore not surprising that we find a high degree of variability in individuals’ ability and the strategies used and declines in normal aging [1], in neurodegenerative disease [2] and with vestibular dysfunction [3,4]. These strategies have been broadly defined as route strategies and orientation strategies [5,6]. The difference between these two strategies is demonstrated in Figure 1.

**Figure 1.**
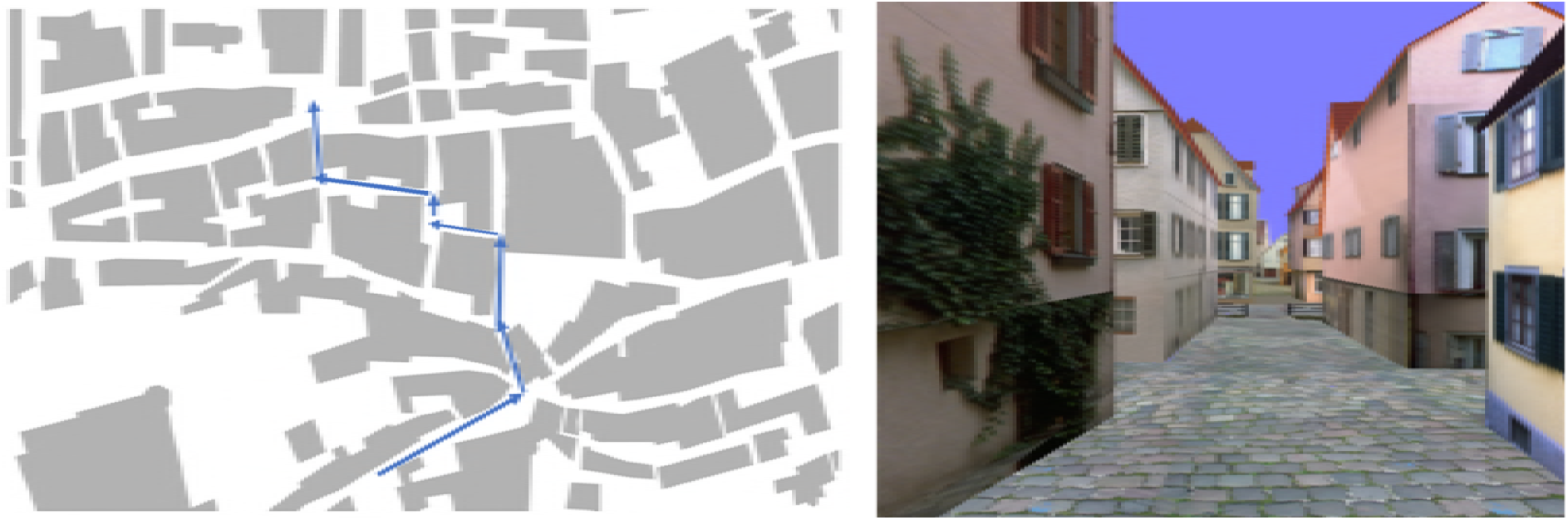
Illustration of the two main wayfinding strategy types. **Left:** the **orientation strategy**, in which individuals use spatial relations and global reference points such as the sun or cardinal directions to navigate. Individuals that use this strategy often report having a 2D map of the environment in their head. An example orientation strategy statement: “*I keep track of the direction (north, south, east or west) in which I was going*”. **Right**: the **route strategy**, where individuals make navigation decisions based upon information in their immediate environment from an egocentric viewpoint. They may know to turn left at the building with ivy on the wall. An example route strategy statement: “*Before starting, I ask for directions telling me whether to turn right or left at particular streets or landmarks*“. Images from Virtual Tuebingen, based on [7,8]

Route strategies, also called “path” or “response-based” strategies involve wayfinding by learning a route from a starting point to a destination as a sequence of instructions based on local orientation points or landmarks cues [1,2,5]. The route strategy requires only an egocentric frame of reference or a path view of the environment [1], in other words the viewpoint that is always present when we navigate through the environment. Knowledge from this path can then be used to assemble the positions into a network of internal representations of landmarks, routes and other positional information. This network of information; often described as a “cognitive map” [6,9] of the environment, provides us with survey knowledge, i.e. an integrated understanding of the spatial layout of the environment, including relative distances between objects. The orientation strategy makes use of this conglomerate of information; global reference points, e.g. the position of the sun, distances or the cardinal directions [5,10,11] are used to navigate the environment. This strategy is thought to rely on an allocentric frame of reference, a bird’s eye perspective or a map-like internal representation of the environment, representations that are independent of one’s current position and orientation in space [9].

Both orientation and route strategies have their advantages and disadvantages. The orientation strategy, when used correctly, allows individuals to take detours or find the correct path when approaching a known intersection from another direction [9,12]. The route strategy is computationally less expensive and therefore likely faster, but individuals are not flexible in finding their way [13]. This suggests that persons that use the orientation strategy have superior wayfinding ability, although individuals may be the most proficient if they are able to combine techniques from both strategies [14].

Neurological disorders can selectively affect wayfinding strategies. Disorientation and impaired wayfinding are often the first signs of senile dementia and Alzheimer’s disease [2,4]. The hippocampus and surrounding medial temporal lobe (MTL) are some of the first brain regions affected by Alzheimer’s disease and dementia [15]. The MTL plays a key role in memory, in particular spatial memory. It is thought to provide important computations for map-based or allocentric navigation [16,17]; computations specific to the orientation strategy. Vestibular dysfunction, the partial or complete loss of function in the vestibular organ or central pathways, is a less well known, but widespread neurological disorder. In an ongoing cross-sectional survey in the United States, vestibular dysfunction, measured by the presence of vertigo/dizziness, was present in over half of the individuals over the age of 40 [18]. Vestibular dysfunction also leads to a decrease in hippocampal size and an associated decrease in navigational ability [3,19,20]. This suggests that Alzheimer’s disease, dementia and vestibular dysfunction may specifically impair the ability for persons to use the orientation strategy. Alternatively, recent evidence suggests that vestibular dysfunction strongly influences cognitive function and navigation in general [3,18,21,22]. It was associated with a decrease in cognitive function equivalent to adding five years of age [18], suggesting that vestibular dysfunction may affect both navigation strategies.

Although we have started to understand the factors that affect strategy use in individuals, there is also much we do not know. One consistent factor that influences wayfinding is gender. Men appear to consistently prefer orientation strategies and generally have superior performance [5,10,23]. Unfortunately, most of the research on wayfinding has either come from small populations [11] or samples with a restricted range of sociodemographic and biological characteristics, primarily college undergraduates [5,10]. We therefore conducted the current study in order to examine the sociodemographic and vestibular components of wayfinding ability in a representative cross-sectional sample of the German population. The three objectives of the study were (1) to investigate the frequency and determinants of wayfinding strategies in a population-based representative sample (2) to test whether vestibular function affects only the orientation strategy or both wayfinding strategies and (3) to examine the frequency of combined scores in the population and how sociodemographic factors influence general wayfinding ability as measured using both strategy scores.

## Methods

### Sample

The data were collected through a computer-assisted telephone interview (CATI-interview) with trained interviewer, as part of an omnibus survey performed by the market research Kantar Health (http://www.kantarhealth.com/). To collect a cross-sectional representative sample from the German-speaking population, the Infratest telephone master sample (ITMS) was designed according to the consortium of German market research institutes (Arbeitsgemeinschaft deutscher Marktforschungsinstitute, ADM-Design) [24]. Participants were recruited if they had a minimum age of 18 and a landline; constituting 90% of all private households in Germany. Participants gave oral informed consent before the questionnaire was administered, in accordance with the Declaration of Helsinki. The telephone numbers were based on the official German telephone registry, stratified according to administrative districts and community sizes. Additional numbers were generated by random selection of the last two digits of telephone numbers. Finally, telephone numbers were randomly selected at the community level. This three-stage sampling design is thought to ensure an unbiased sample selection that excludes clustering effects and allows a random selection of defined targets. For example, the lifetime prevalence of 25-30% expected for vertigo/dizziness was estimated with a precision of 2.7% from a sample size of 1,000 participants using the same sampling design [25,26]. Data collection ran December 11th, 12th and 16th 2015.

### Measures

The Wayfinding Scale [5], was used to determine relevant predictors of wayfinding strategy types and wayfinding ability. The original scale comprises a total of 14 items; nine items for the orientation strategy, with a maximum score of 45, and five items for the route strategy, with a maximum score of 25. For every item on the original scale, there is a 5-point-likert answering scale, where 1 means *not at all typical for me* and 5 *very typical for me*. For the purpose of this study we deleted one item concerning orientation strategy (“I refer to a published road map when I drive”) because of the decreased use of written maps and increase use of mobile devices for navigation. Additionally, an item from 2002 modified International Wayfinding Scale [10] (“I found maps of the building or complex, with an arrow pointing to my present location, to be very helpful”) replaced one item concerning the route strategy (“Before starting, I ask for a hand-drawn map of the area”) for similar reasons. Consequently, the Wayfinding Scale used had 13 items instead of 14 items and a maximum attainable score of 40 instead of 45 for orientation strategy. The questions used in our Wayfinding Scale can be found in the supporting information (S1 File). For further analysis, the individual sum scores from each strategy were scaled to 100 to compare the individual wayfinding ability among the two strategies and to determine which strategy was preferred.

The original scale was adapted for German by multiple iterations of a translation-retranslation-procedure by a team of German and English native-speakers [27]. Because the Wayfinding Scale has been used several times [5,10,28] in different countries [29] and associated with real-world navigational tasks [11], it is regarded as valid and transferrable for the current study. Sociodemographic characteristics included age, sex, education, household income, and a nationally defined regional urbanization metric. Age was stratified into four brackets (18-35, 36-55, 56-70, 71-96). The division at age of 55 was near to the median age and allowed to include the thesis of hormonal regulation [30,31]. Education was measured by the highest level of academic achievement. Level of education was then grouped according to the International Standard Classification of Education (ISCED 97) [32,33] into primary/lower secondary, secondary/non-tertiary, upper secondary and tertiary education. The 15 individuals still in school were grouped with the participants who were in secondary, non-tertiary education. Education and household net income were stratified into quartiles. Two items assessed vertigo and balance: “Did you experience moderate or severe dizziness or vertigo during the last 12 months? (rotational vertigo, staggering vertigo, imbalance) and “How good is your sense of balance compared to other people your age?”. The town sizes are shown according to BIK regions, which is a national regional classification system established by the market research institute BIK Aschpurwis + Behrens GmbH, comparable to the Metropolitan Statistical Areas (MSA) in the USA [24]. BIK systematics better express the structural features of today’s city regions than the Boustedt method (or the current political town size classes in the new federal states). Existing municipalities in Germany are defined as BIK regions according to the number of inhabitants of a catchment area and the size and intensity of commuter links [34,35]. The BIK-regions can be seen as a measure of regional urbanization, classified by the number of inhabitants per region.

### Statistical analysis

Categorical variables were summarized by frequencies and percentages. Continuous variables were summarized by mean and standard deviation.

To adjust for the effect of over- or under-representation of specific person groups, e.g. the over-representation of middle aged and upper-/middle-class participants, the sample was weighted by federal state, regional division system, age, gender, occupation, education and the number of individuals living in a household. Thus, each target person was fitted with an individual weighting coefficient. The weighting coefficients summed to the sample size, while the mean value of the weights across the sample was equal to one. Individual weightings ranged from 0.27 to 12.69. A weighting of less than one reduced the effect of an over-represented person and a weight greater than one was meant to adjust the influence of participants that were underrepresented in the sample [32,36,37]. If not stated otherwise, we present the weighted results.

To investigate the determinants of wayfinding, we applied separate linear models for both orientation and route scales as outcomes using the Wayfinding Scale score values (S2 Table). Regression diagnostics for all models included tests for multicollinearity using the variance-inflation-factor and residual-plots, Breusch-Pagan-Screening-Test for heteroscedasticity and Kolmogorov-Smirnov-test for normal distribution as well as further residual diagnostic like cook-distance for outliers [38–41]. Since regression diagnostic of the route scores indicated heteroscedasticity and a variance-stabilizing log-linear-transformation of the scores did not improve goodness-of-fit, regression analysis with heteroscedasticity-consistent standard errors was used with a heteroscedasticity consistent covariance matrix (HCCM 0) based on the Huber-White-Eicker weighting procedure for standard errors. This method is the procedure of choice in this situation as the sample size was large enough (n>250), and because it allows for easy interpretation [42].

Using separate models for the route and the orientation strategy neglects the fact that the two scores come from the same individual. Some individuals may score high in both route and orientation strategies, allowing them to flexibly adapt their wayfinding strategy to meet the needs of the situation. Similarly, individuals may score low on both strategies, suggesting that they have a difficult time successfully wayfinding in any situation. To investigate the combined outcome, we categorized individuals into four distinct classes based on their medians of both strategies (76 points on route scale, 60 points on orientation scale). Because our sample is large and representative, the median cut-off values used here can be applied to other studies with smaller and less representative samples.

An individual that scored at least 76 points on the route scale, but below 60 points on the orientation scale was classified as a route strategist. An individual that scored 60 points or more on the orientation scale, but less than 76 points on the route scale was classified as an orientation strategist. An individual scoring at least 76 points on route scale and at least 60 points on orientation scale was categorized as a “flexible” strategist. A person that scored less than 76 points on route scale and less than 60 on orientation was defined as an “undetermined” strategist. We then used this categorized outcome for a multinomial regression [41,43].

All data analysis was carried out using SPSS software (IBM SPSS Statistics for Windows, Version 23.0. Armonk, NY: IBM Corp) and the IBM R-Essentials [44] using R software, version 3.1.0 [45].

## Results

One-thousand three participants, aged 18-96 agreed to participate in the survey. All participants with missing values were excluded from the statistical analyses. The final sample included 783 participants; 52.7% were women and the mean age was 47.9 years (SD = 17.9). Prevalence of vertigo/dizziness was 24.2%. Table 1 shows the sociodemographic characteristics of all participants that were included in the regression analysis, separated by strategy. As described in the Mehods, all data is weighted according to the frequency of the current population.

**Table 1.**
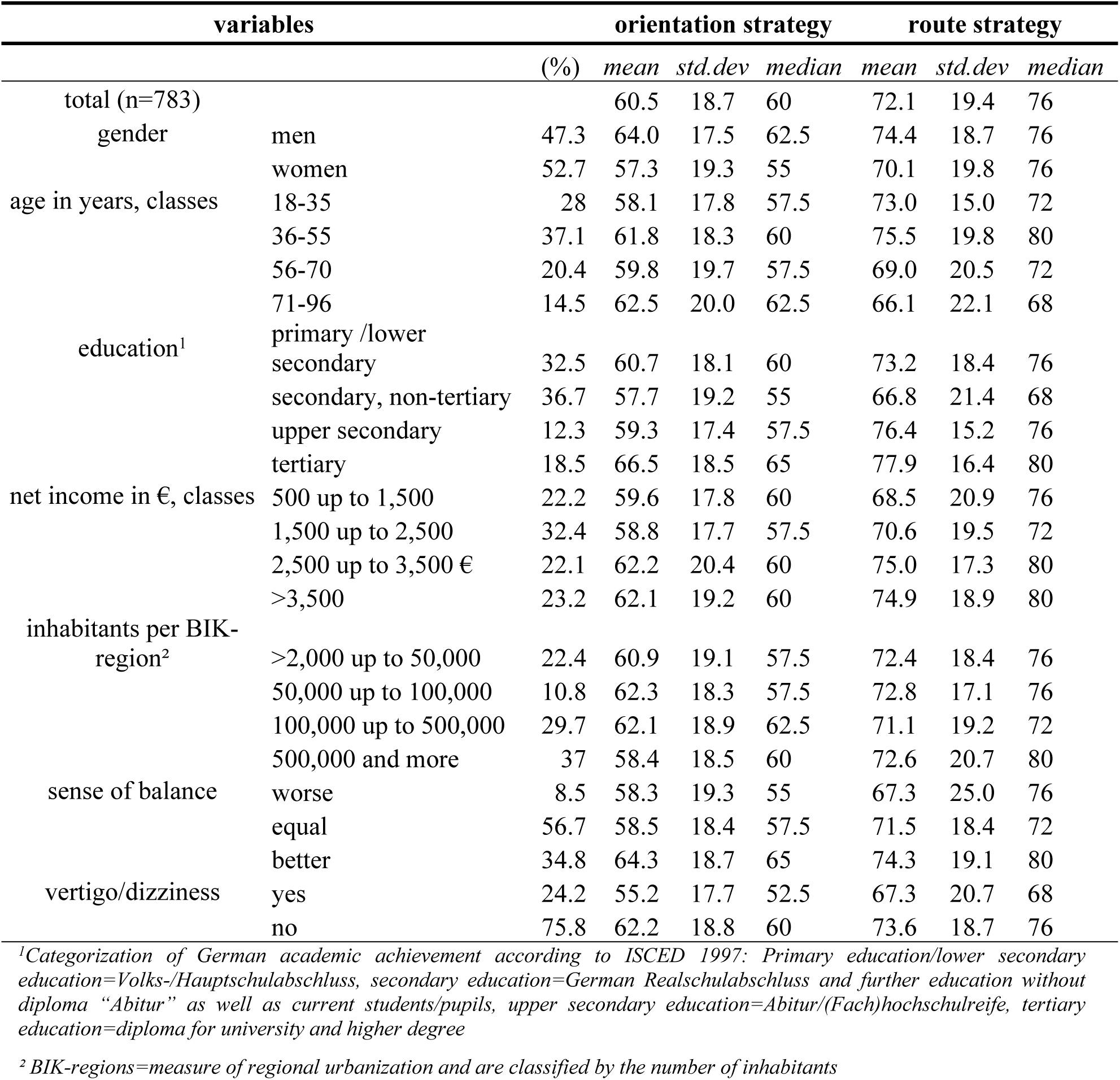
Scores for the orientation and route strategy from the Wayfinding Scale, stratified by each sociodemographic class, including vertigo and balance (n = 783). The outcome scores are scaled to 100 and individually weighted according to the frequency of the current population

## Individual wayfinding strategies

Two linear regressions were performed to examine the effect of sociodemographic determinants and vestibular performance, on the orientation and the route strategy respectively. The results of each of these regressions are presented in Table 2.

**Table 2.** Results of the multiple regression for each strategy separately (n = 783). Beta-coefficients (ß), Confidence Intervals and p-values for the regression coefficients for orientation strategy and route strategy (with heteroscedasticity-robust standard errors – see Methods), and individually weighted according to frequency of the current population.

In the orientation strategy gender differences across the entire population were in accordance with what has been seen in other studies with a more limited age and educational stratification. Men reported significantly higher orientation scores than women by almost 5 points. The influence of estrogen was examined by comparing postmenopausal women in both age categories above the age of 55 to younger women, with the expectation that women above 55 years of age have less estrogen and therefore higher orientation scores. However, we found no evidence that post-menopausal women reported higher orientation scores than pre-menopausal women.

In contrast to our expectations, older participants scored higher on the orientation strategy than younger participants. A person aged 71 or older reported an orientation strategy score that was on average 7 points higher than someone under the age of 36. Educational level and regional urbanization influenced orientation strategy scores in opposite directions. Increasing educational achievement lead to significantly higher orientation scores. Participants residing in urban areas with over 500,000 inhabitants had significantly lower orientation scores than residents in areas with up to 50,000 inhabitants.

Our second objective involved understanding the relationship between wayfinding and vestibular function. Both measures of vestibular performance had significant effects on the orientation strategy. Persons with vertigo in the last 12 months had reduced orientation scores of over 6 points. Correspondingly, participants with a good sense of balance had significantly higher scores on the orientation strategy scale.

In accordance with previous research, the route strategy showed much less stratified effects than the orientation strategy. There was no significant effect of age or gender on route strategy scores. However, two interesting and significant effects were found that were also seen in the orientation strategy. First, participants with higher educational achievement also reported increased scores on the route strategy. Second, the presence of vertigo in the last 12 months was associated with a 5-point decrease in route strategy scores compared to participants without vertigo.

In summary, the relevant sociodemographic determinants for wayfinding proved to be gender, age, regional urbanization and education. Income was the only factor measured that did not significantly influence wayfinding scores.

## Combined wayfinding strategies

The Wayfinding Strategy Scale provides two independent scores for each participant; one for the orientation strategy and one for the route strategy. If analyzed separately, as has always been done previously, these scores do not show combined effects across both strategies. Most studies agree, though, that superior wayfinding involves the ability to switch between different strategies for flexible and fast adaptation to the situation at hand [1]. We therefore chose a novel approach to analyze the Wayfinding Strategy Scale, taking advantage of our representative sample. We grouped our participants into four groups using a median split and examined the sociodemographic determinants of combined strategies using a multinomial regression model and including all predictors from the linear regression models. Interestingly, the majority of individuals reported using either both strategies or neither strategy, instead of preferring one strategy over the other: 1) 30.7% were **undetermined strategists** that scored below the median in both strategies, 2) 18.5% were **route strategists** that scored above the median only in the route strategy, 3) 15.5% were **orientation strategists** that only scored above the median in the orientation strategy, and 4) 35.3% were **flexible strategists** that scored above the median for both strategies.

The multinomial model is highly complex with all possible combinations of differences between groups. However, the odds ratios provide a useful way of interpreting the results of the analysis. An odds ratio greater than 1 means that there is a positive effect of that sociodemographic factor grouping to use a specific strategy compared to the reference grouping and strategy, whereas less than 1 means there is a negative effect. The multinomial model confirmed the results from the linear regression concerning the orientation strategy vs. the route strategy and will therefore not be reported here (for the full model see S3 Table). Instead, we focused here on flexible strategists, who have the ability to use both strategies of wayfinding and should therefore be superior navigators and compared their odds ratios to the undetermined strategists and the orientation strategists (Table 3).

**Table 3.** The odds of being a flexible strategist (n = 783). Odds ratios (OR), Confidence Intervals (CI) and p-values (p) from the multinomial regression model for the odds of flexible vs. undetermined and flexible vs. orientation strategies. Outcomes were individually weighted according to frequency distribution of the current population. OR >1 means it is more likely to be part of the group of interest.

Comparing the flexible strategy to the undetermined strategy, a flexible strategist had greater odds of being male, having a high education level, and being older in age (in reference to the youngest age group of 18-35). Comparing the flexible strategy to the orientation strategy, a flexible strategist had greater odds of being male, having a high education level, and living in a lower-density urban area with 100,000 up to 500,000 inhabitants per region. Men were more likely to use a flexible strategy than an undetermined strategy compared to women but did not have significantly higher odds of being a flexible vs. an orientation strategist. Persons in the age group 36-55 yrs. and 71-96 yrs. had significantly greater odds of being a flexible strategist than being an undetermined strategist. However, the age group 71-96 did not have greater odds of being a flexible strategist compared to an orientation strategist.

In summary, older males, with a higher education and living in less urbanized areas tended to report using both the orientation and the route strategy, although gender and age effects were similar between the flexible strategy as well as the orientation strategy. In addition, the presence of vertigo, and being in the youngest age group (18-35 years) also reduced the odds of using a flexible strategy for wayfinding.

## Discussion

Using the wayfinding strategy scale, we examined how sociodemographic measures influence whether a person tends to follow a route or develop a map of the environment. Persons living in less urban regions, having higher education, being male or over the age of 35 were more likely to report using a map-based wayfinding strategy (the orientation strategy). Being younger, being female or living in more urban areas were indicative of lower scores in the orientation strategy. The presence of vertigo/dizziness in the last 12 months decreased scores for both wayfinding strategies, implying that vestibular problems impair general wayfinding ability. To look at combined effects across wayfinding strategies, we grouped persons with high scores in both strategies as flexible strategists and persons with low scores in both strategies as undetermined strategists. Individuals tended to use both strategies if they were over 35 years old, well-educated and living in less urban areas. Our results provide new insights into how environment, education and behavior affect how humans navigate across an entire adult lifespan.

One of the factors that most consistently affects wayfinding is gender. The fact that men report higher orientation strategy scores than women has been shown in young adults [5,11,29]; we demonstrate the same trend across all age groups. Men also have higher scores for both the route and orientation strategy, suggesting they can flexibly choose what navigation strategy to use. This may explain the overall and a task-related advantage in navigational ability in real and simulated environments, albeit in only about half of the cases (49.28%) [23]. Gender differences already exist in childhood, but it is not clear to what extent biological and sex-typed experiential factors interact [10,55–57] to produce this effect and if they are consistent throughout lifetime [9].

Because we measured a large range of ages, we were able to show that age had a large effect on wayfinding scores. Older participants showed a stronger reliance on orientation strategy and overall higher scores than younger participants. In the original Wayfinding Scale study, older participants also tended to report using the orientation strategy, which was attributed to a growing experience in older persons [5]. Most of their participants, however, were between the ages of 18-35, which encompasses our youngest age group. The high scores reported by the oldest age group in our study represent novel findings, that are not entirely consistent with the literature. Behavioral experiments on real or virtual navigation report that older persons have difficulties switching between strategies [58], forming and using a cognitive map [47] and that they prefer an egocentric strategy [47] for navigation, such as the route strategy. Other studies have shown spatial memory deficits among older persons in mental rotation tests [4], and virtual learning/wayfinding performances, typically measured by errors, distance and/or speed [48,49,59]. Navigational self-reports from older participants tend to inflate their actual navigational ability [49] (but see [50]), suggesting that the high scores in the oldest age groups in our study may result from inaccurate reporting. However, the age group 36-55 was more likely to flexibly use both strategies than the youngest age group, and we would not expect the reporting biases to already be present here. Future studies that examine self-reported wayfinding preferences and behavior within the same individual would disentangle these effects.

In general, the average scores for both wayfinding strategies were higher across our sample than in previous studies. Although this could suggest a general increase in reporting over time, we attribute it to the greater age diversity in our sample, in particular the higher number of older participants. Previous studies have used smaller sample sizes or narrower age ranges [5,10,11,29,53], therefore, we believe the higher scores are more representative of wayfinding across the population.

The strongest positive influence on wayfinding ability comes from education. This could simply be a result of a self-report education bias, where participants with higher education had a better self-assessment [51]. However, behavioral research on visuospatial attention and cognition demonstrate a relationship between higher education and better spatial ability [52,55]. Higher education monotonically increases the probability of a person using the orientation strategy and is indicative of a person being a flexible strategist. Whether the educational effect on wayfinding strategy is a result of improved spatial ability remains to be seen.

Participants living in urban areas reported lower scores on orientation strategy, emphasizing the idea that the geographical topography of the environment influences the wayfinding strategy used. Previous research has shown that both men and women are more likely to use cardinal directions when giving directions if they came from places laid out in a grid-like pattern [53]. In cities, the omnipresence of signs and buildings makes it impossible or unnecessary to orientate via distances and cardinal directions as in the orientation strategy.

Previous studies have shown an age-gender interaction for the orientation strategy, where men show a greater increase in the orientation strategy scores with increasing age [5]. Here instead, we found an age-gender-interaction for route strategy, where younger persons, especially younger men, rely more on the route strategy than older persons. Recent work suggests that the use of mobile GPS devices for navigation activates less of the brain, in particular in the hippocampus, and area thought to be important for the orientation strategy [60] and also leads to increased errors in navigation [61]. This decrease in the use of the orientation strategy in young men could be the first population-based evidence of behavioral changes resulting from increased GPS usage, particularly in the younger generation. We are currently specifically examining the effect of GPS-use on the choice of wayfinding strategy.

Participants with vertigo or dizziness, even if they do not have a clear vestibular pathology, have a disadvantage in wayfinding. Similarly, patients with vestibular loss have difficulties in spatial memory [62–65] and wayfinding tasks [3,19,20] as well as reduced hippocampal volume. The hippocampus is thought to be important for the allocentric navigation [9,67] that forms the basis of the orientation strategy. However, participants with vertigo had low scores on both the orientation and the route strategy, which cannot totally confirm the connection between vestibular dysfunction, allocentric navigation and the hippocampus. Our results emphasizes that vestibular input is an important source of information for spatial memory and efficient wayfinding [17,66] for both wayfinding strategies, and supports the theory that vertigo and dizziness has a more generalized effect on cognition [18], more than a specific effect on spatial memory. We are aware that we included a broad definition of vertigo/dizziness. However, the vertigo symptoms were assessed by standardized questions derived from previous studies [54] and the prevalence of vertigo corresponded to recent findings [26].

Similar to previous studies using the Wayfinding Scale, the route strategy showed higher scores overall than the orientation strategy, emphasizing the idea that route strategy is less computationally expensive, and therefore less challenging than orientation strategy [5,11,28,67]. To examine the ability to switch between different strategies we grouped our sample into four groups, the predominant orientation strategist, the route strategists and our two novel groups, the flexible strategists that use both strategies, and undetermined strategists that do not appear to use either strategy. Males with a high education and living in more rural areas are more likely to flexibly use both strategies, in line with behavioral evidence for a male and educational advantage in spatial abilities [52]. Having vertigo and being in the youngest age group was indicative of not using either strategy, confirming the effect vestibular dysfunction and GPS use on general navigational ability.

## Conclusion

Our study is the first to show the strong influence of all sociodemographic factors except income in the choice of wayfinding strategy in a representative sample of the population. We specifically demonstrate the detrimental effect of vertigo and dizziness on wayfinding ability, and a potential change in the way persons (especially young persons) navigate, as a result of the increased use of mobile GPS devices. Plausible mechanisms for these effects may involve orientation-specific brain areas and effects of vestibular input on cognition [3,19,20]. The scores acquired can be used for comparisons in future studies with smaller sample sizes. Longitudinal studies and experiments involving specific navigational paradigms are needed to understand the underlying mechanisms for the sociodemographic effects found here.

## Supporting information

**S1 File.** The wayfinding strategy scale as used in the questionnaire

**S2 Table.** Alternative linear regression models, including lin-log regression and age in yrs. (metric)

**S3 Table**. Multinomial linear regression models for all possible odds concerning the undetermined, route, orientation, and flexible strategy.

**Data availability**. Data for all studies are available at https://web.gin.g-node.org/.

**Code availability**. SPSS Syntax-file is available at https://web.gin.g-node.org/. The file includes preparation and all relevant analysis for this paper.

## Acknowledgements

This project was supported by funds from the German Federal Ministry of Education and Research under the Grant Code 01EO1401. The authors bear full responsibility for the content of this publication.

